# Gag ubiquitination facilitates ALIX-mediated lentiviral RNA packaging, counteracted by USP8

**DOI:** 10.1101/2025.05.27.656518

**Authors:** Kenta Hagiuda, Yuya Himi, Masayuki Komada, Toshiaki Fukushima

## Abstract

Lentiviral Gag protein plays an integral role in viral assembly. In host cells, Gag multimerizes, recruits host budding factors, and buds from the plasma membrane. Additionally, it binds to viral genomic RNA (vRNA) and facilitates its packaging into viral particles. However, the mechanisms regulating Gag-vRNA interactions remain poorly understood. Previous studies have suggested that Gag ubiquitination influences viral infectivity. In this study, we elucidated a new mechanism by which Gag-vRNA interactions are regulated via Gag ubiquitination. Using an HIV-1-derived lentiviral vector system, we found that ubiquitin-specific protease 8 (USP8), a deubiquitinating enzyme involved in antiviral responses, reduced the vRNA content in virions through its catalytic activity. USP8 deubiquitinates Gag, and mutations at the Gag ubiquitination sites similarly impair vRNA packaging. We hypothesized that Gag ubiquitination enhances its interaction with RNA-binding proteins, thereby facilitating Gag-vRNA binding. We focused on ALG-2-interacting protein X (ALIX), which interacts with Gag, ubiquitin, and RNA. Mechanistic analyses revealed that Gag ubiquitination promotes its interaction with ALIX, which in turn binds to vRNA. In ALIX knockout (KO) cells, both Gag-vRNA binding and vRNA packaging were reduced, confirming the role of ALIX in this process. Furthermore, the deubiquitinating activity of USP8 suppressed Gag-vRNA binding, and this effect was abolished by mutation of the Gag ubiquitination site or ALIX KO. Collectively, these results reveal a new regulatory mechanism for lentiviral assembly, in which Gag ubiquitination facilitates ALIX-mediated vRNA packaging and is counteracted by USP8.

**Significance Statement:** Lentiviral Gag-vRNA binding and vRNA packaging are essential for viral particle formation; however, their regulation remains unclear. We identified a USP8-Gag-ALIX axis that controls this process. USP8, a host deubiquitinating enzyme, suppresses Gag-vRNA binding by removing ubiquitin from Gag. Gag ubiquitination promotes ALIX interaction, which facilitates Gag-vRNA binding. Although ALIX is known to play a role in viral budding, we uncovered a new role for ALIX in vRNA recruitment. These findings advance our understanding of lentiviral assembly and identify USP8 and ALIX as potential targets for antiviral therapy. They may also improve lentiviral vector production by providing a molecular basis for reducing transgene-deficient particles and enhancing vector quality.

## Introduction

Lentiviruses are a subgroup of retroviruses that cause chronic infections and fatal diseases in mammals. Lentiviral particles comprise viral genomic RNAs (vRNAs), viral proteins, and a lipid envelope. The viral Gag protein plays a central role in the packaging of vRNAs into viral particles (1). In the host cytoplasm, Gag binds to vRNA, and the resulting complex is transported to the plasma membrane (2). Gag molecules oligomerize to form large ribonucleoprotein complexes containing two copies of vRNA and thousands of Gag molecules (2, 3). Gag also recruits components of the endosomal sorting complexes required for transport (ESCRT), which facilitates membrane scission and viral budding (4). During and after budding, Gag is proteolytically processed and reorganized to generate mature virions (5). Each Gag domain plays a distinct and essential role in this process (6). The matrix (MA) domain interacts with the plasma membrane, the capsid (CA) domain mediates Gag multimerization through homotypic interactions, the nucleocapsid (NC) domain binds to the vRNA, and the p6 domain recruits the ESCRT components.

Although viral assembly is an organized process, it is not error-free. Gag can form virus-like particles (VLPs) in the absence of vRNA (7), indicating that non-infectious vRNA-deficient particles are inevitably produced. To compensate for this inefficiency, lentiviruses may have evolved mechanisms to enhance the fidelity of vRNA packaging, whereas host cells may apply counteracting pressures to impair it. In practice, improving vRNA packaging efficiency remains a major challenge for lentiviral vector production (8). However, the molecular mechanisms that regulate vRNA packaging remain poorly understood.

One possible regulatory mechanism involves the post-translational modification of Gag. Gag is ubiquitinated in host cells (9, 10). Mutations at ubiquitination sites reduce viral infectivity (11), whereas overexpression of the responsible E3 ligases, neural precursor cells expressing developmentally downregulated protein 4 (Nedd4)-1 and -2, enhances viral budding (12–16). These findings suggest that lentiviruses exploit the host ubiquitination machinery to promote their assembly. However, whether Gag ubiquitination is reversible via host deubiquitinating enzymes (DUBs) and how it modulates viral assembly remain unclear.

We investigated the cellular functions of ubiquitin-specific protease 8 (USP8), a DUB known to regulate membrane trafficking, growth factor and stress signaling, and immune and inflammatory responses (17–23). In this study, we generated lentiviral vectors encoding USP8 as research tools and unexpectedly found that its deubiquitinating activity negatively regulated viral titers. We identified USP8 as a DUB that targets Gag and demonstrated that USP8 acts as a host factor that suppresses lentiviral infectivity. Further mechanistic analysis revealed that ubiquitinated Gag interacts with the ESCRT accessory protein ALG-2-interacting protein X (ALIX), which in turn facilitates the Gag-vRNA interaction. These findings reveal a host-mediated regulatory mechanism that controls vRNA packaging into lentiviral particles.

## Results

### USP8 negatively regulates lentiviral vRNA packaging through its deubiquitinating activity

We generated lentiviral vectors encoding USP8 using a standard packaging system in which viral particles were produced in HEK293T cells transiently expressing viral components. Lentiviral vectors encoding a catalytically inactive mutant of USP8 (USP8^C786A^) exhibited higher titers than those encoding the wild-type USP8 (Fig. 1A), indicating that the deubiquitinating activity of USP8 negatively regulates viral titers.

**Figure 1.**
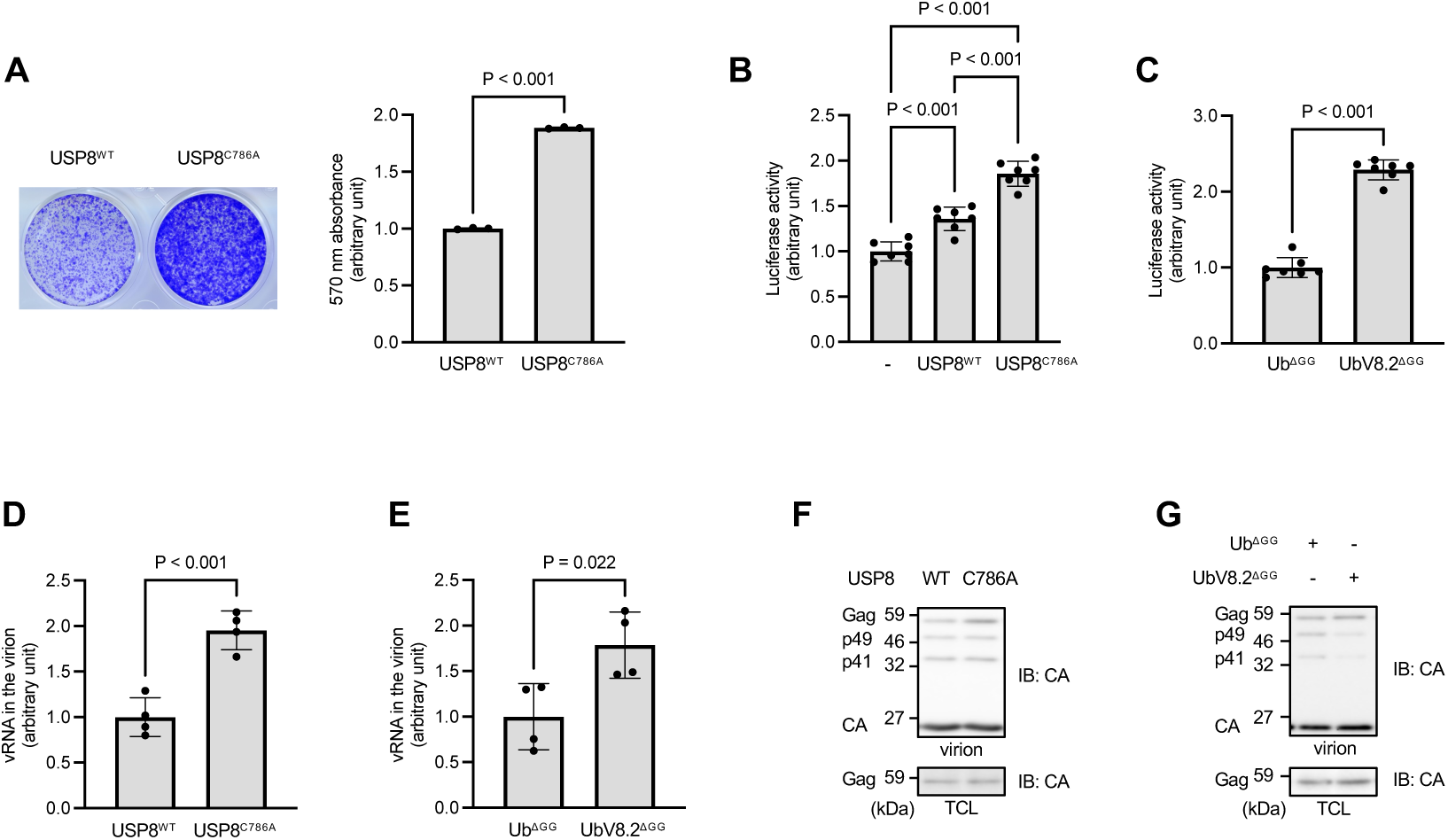
USP8 negatively regulates lentiviral vRNA packaging through its deubiquitinating activity. (A) Lentiviral vectors for the co-expression of either wild-type or catalytically inactive (C786A) USP8 and puromycin resistant genes were produced in HEK293T cells and used to infect HeLa cells. Infectivity was assessed by puromycin selection, followed by crystal violet staining. Left: representative images of stained dishes. Right: absorbance at 570 nm measured from the dye extracts. (B) Lentiviral vectors encoding a luciferase reporter were produced in HEK293T cells transiently transfected with lentivirus plasmids and expression plasmids (mock, USP8^WT^, or USP8^C786A^) and used to infect HeLa cells. Infectivity was quantified by luciferase assay. (C) Experiments were performed as in B, except that the indicated ubiquitin-related plasmids were used instead of USP8 expression plasmids. Ub^ΔGG^: a ubiquitin mutant lacking the C-terminal diglycine motif (ΔGG), rendering it incapable of serving as a donor in ubiquitination reactions. UbV8.2^ΔGG^: a USP8-specific inhibitory ubiquitin mutant also lacking the ΔGG motif. (D-G) Viral particles were pelleted by ultracentrifugation from the culture media of HEK293T cells prepared in B and C. The vRNA content in the viral fractions was quantified using RT-qPCR (D, E). The amount of Gag and its processed forms in the viral fractions and total cell lysates (TCLs) were analyzed by immunoblotting (F, G). Graphs show means ± SD from three (A), seven (B, C), and four biological replicates (D, E).

In this study, USP8 or its mutant was overexpressed in both virus-producing and target cells. To specifically assess the role of USP8 deubiquitinating activity in virus-producing cells, we produced lentiviral particles encoding NanoLuc (Promega) in HEK293T cells expressing either wild-type USP8 or USP8^C786A^ and used the resulting media to infect target cells. Infectivity was quantified by measuring luciferase activity in cell lysates post-infection. The media from USP8^C786A^-expressing cells showed approximately 35% higher infectivity than the media from wild-type USP8-expressing cells (Fig. 1B). Interestingly, wild-type USP8 expression also increased infectivity by approximately 35% compared to the control, suggesting that USP8 exerts both enzymatic activity-dependent and -independent effects, potentially acting as a scaffold protein to promote infection, whereas its enzymatic activity suppresses viral production. Supporting this notion, the expression of UbV8.2^ΔGG^, a ubiquitin variant that selectively inhibits USP8 (24) (Fig. S1A) increased infectivity by approximately 130% compared to that of the control (Fig. 1C), confirming that the enzymatic activity of USP8 inhibits virus production.

To elucidate the underlying mechanism, we purified virions from the culture media by ultracentrifugation and measured the vRNA and Gag protein content. The vRNA levels in virions increased by approximately 100% in USP8^C786A^-expressing cells compared to those in wild-type USP8 (Fig. 1D) and by approximately 75% in UbV8.2^ΔGG^-expressing cells (Fig. 1E), whereas intracellular vRNA levels remained unchanged (Fig. S1B and S1C). In both cases, the total amount of Gag and its processed forms in the virions were not significantly altered (Fig. 1F and 1G). These results suggest that USP8 negatively regulates vRNA packaging via its deubiquitination activity.

### USP8 deubiquitinates Gag

We speculate that USP8 may act as a DUB for Gag. Co-immunoprecipitation revealed that USP8 interacted with Gag (Fig. 2A), and immunostaining showed co-localization in the cytoplasmic puncta (Fig. 2B). To assess whether USP8 regulates Gag ubiquitination, immunoprecipitation (IP) and immunoblotting (IB) were performed. Among several ubiquitinated species detected as bands and smears in Gag immunoprecipitates, only a ∼60 kDa band remained after SDS denaturation prior to IP (Fig. 2C), probably corresponding to mono-ubiquitinated Gag (unmodified Gag, approximately 55 kDa), consistent with previous reports (9, 10). Gag monoubiquitination was increased by USP8 knockdown (Fig. 2D), decreased by the expression of wild-type USP8 compared to USP8^C786A^ (Fig. 2E) and increased by the expression of UbV8.2^ΔGG^ (Figs. 2F and S2), indicating that USP8 functions as a DUB for Gag.

**Figure 2.**
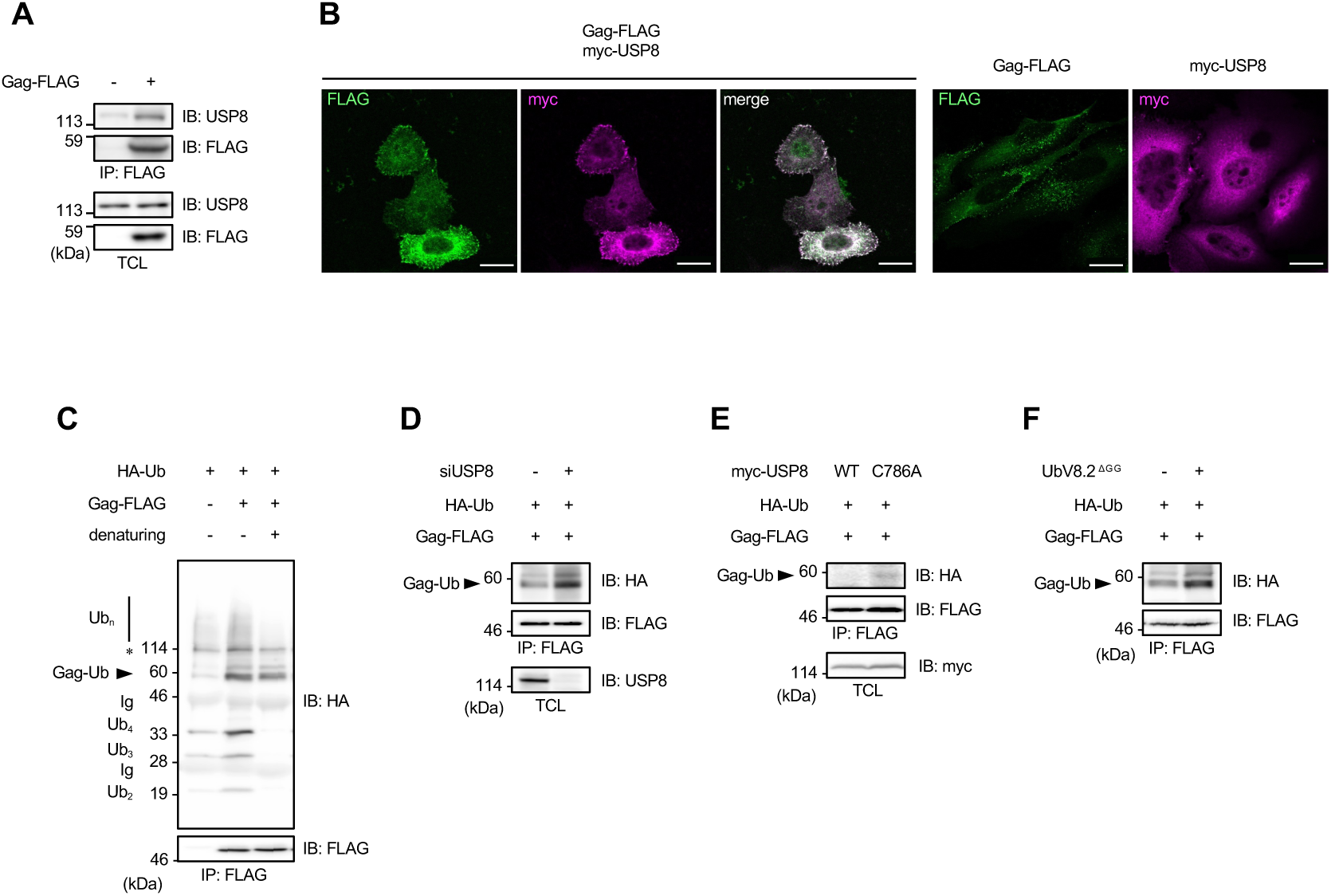
USP8 deubiquitinates Gag. (A) HEK293T cells transfected with Gag expression plasmids were subjected to crosslinking followed by immunoprecipitation (IP). Immunoprecipitates and TCLs were analyzed by immunoblotting (IB). (B) HeLa cells transfected with the indicated expression plasmids were analyzed by immunofluorescence staining. Scale bar, 20 μm. (C-F) HEK293T cells transfected with the indicated expression plasmids or siRNAs were subjected to IP and IB. In C, some samples were denatured with SDS treatment and boiling prior to IP.

### Gag ubiquitination promotes vRNA packaging into lentiviral particles

To examine the role of Gag ubiquitination, we mutated known ubiquitination sites in the NC, spacer peptide 2 (SP2), and p6 domains (11). Two mutants were generated: Gag^14KR^ (all 14 lysine residues were replaced with arginine) and Gag^4KR^ (four lysine residues in SP2 and p6 were replaced) (Fig. 3A). Both mutants showed a marked reduction in vRNA incorporation into the virions (approximately 20% of the wild-type level) (Fig. 3B and 3C) without affecting intracellular vRNA levels (Fig. S3A and S3B). The total levels of Gag and its processed forms in the virions did not decrease (Fig. 3D and 3E). These results indicate that Gag ubiquitination facilitates vRNA packaging into lentiviral particles.

**Figure 3.**
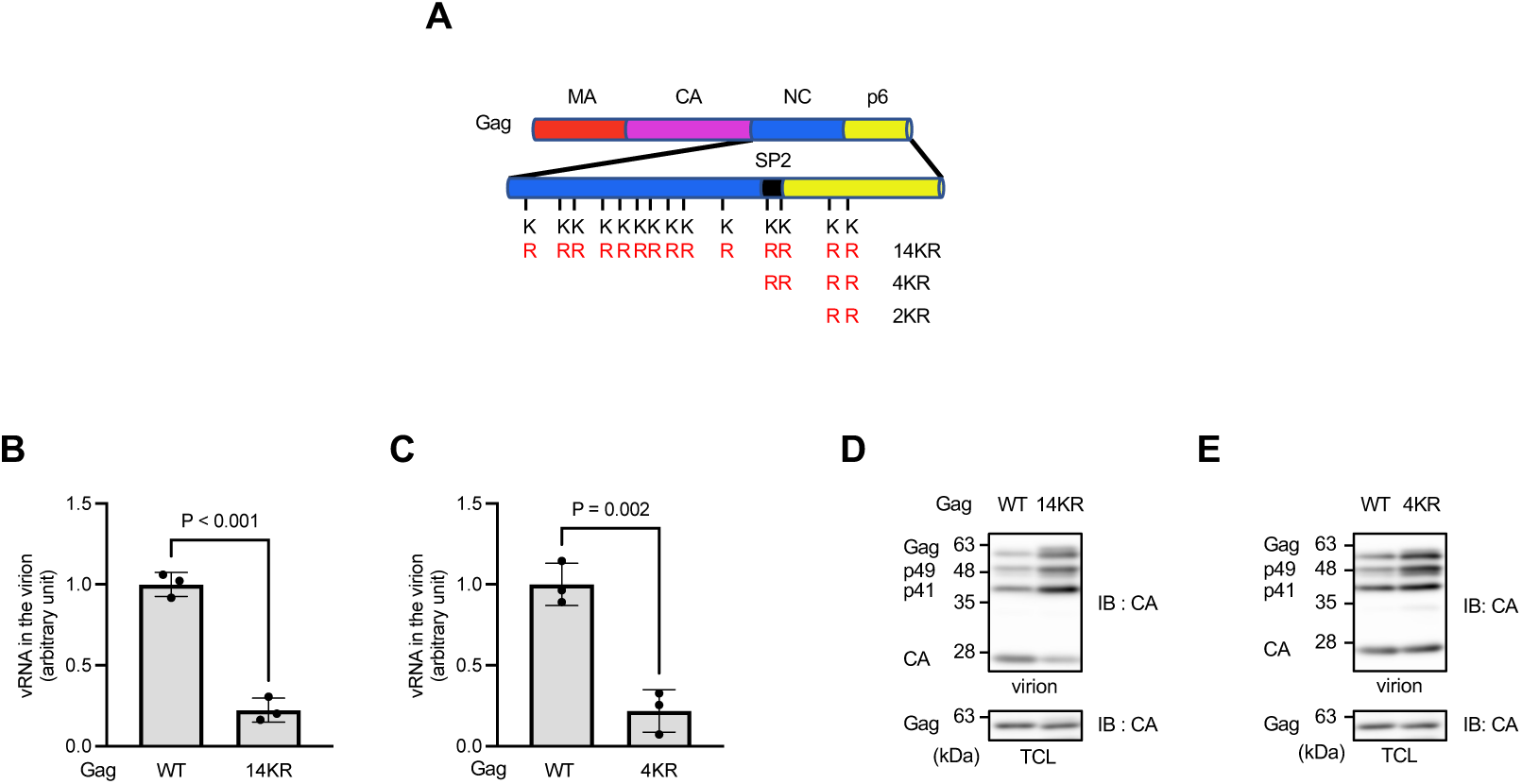
Gag ubiquitination promotes vRNA packaging into lentiviral particles. (A) Schematic of Gag mutants used in this study, in which lysine residues at ubiquitination sites are replaced with arginines. (B-E) Lentiviral vectors encoding a luciferase reporter, with or without Gag ubiquitination site mutations, were produced in HEK293T cells. Viral particles were pelleted from the culture media by ultracentrifugation, and vRNA content in the viral fractions was quantified by RT-qPCR (B, C). The amount of Gag and its processed forms in the viral fractions and TCLs were analyzed by IB (D, E). Graphs show means ± SD from three biological replicates.

### Gag ubiquitination promotes its interaction with ALIX, which binds to vRNA

We hypothesized that Gag ubiquitination enhances its interaction with RNA-binding proteins, thereby facilitating Gag-vRNA binding. A probable candidate is the ESCRT accessory protein ALIX, which contains Bro1 and V domains, as well as a C-terminal proline-rich region (PRR) (Fig. 4A). ALIX interacts with Gag via Bro1–NC and V–p6 interactions (25, 26). The Bro1 domain has been suggested to possess RNA-binding ability (27), whereas the V domain binds to ubiquitin (28, 29).

**Figure 4.**
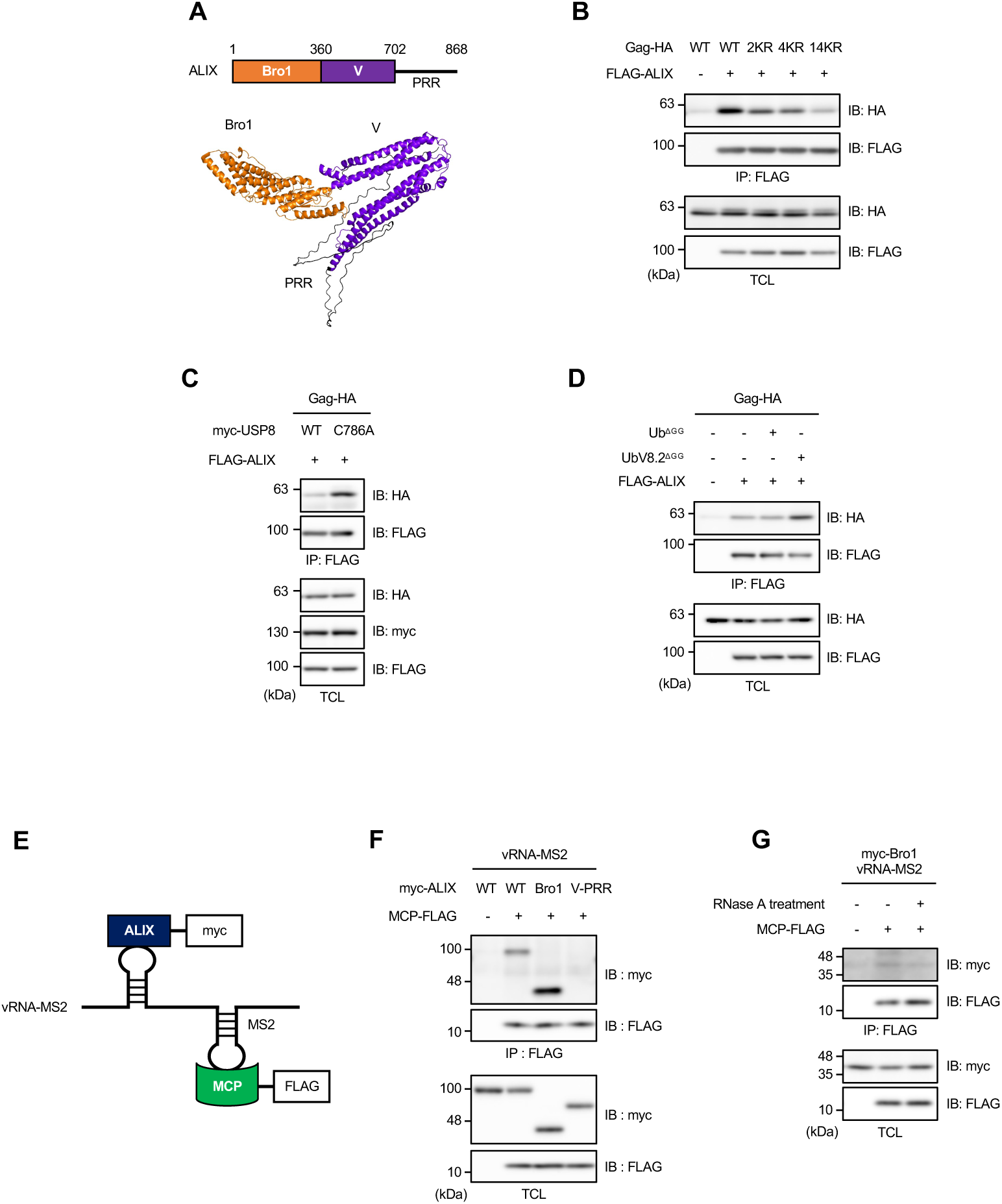
Gag ubiquitination promotes its interaction with ALIX, which binds to vRNA. (A) Domain architecture of ALIX (top) and its 3D structure predicted by Alphafold3 (bottom). (B-D) HEK293T cells were transfected with the indicated expression plasmids. Cell lysates were subjected to IP. Immunoprecipitates and TCLs were analyzed by IB. (E) Schematic of the MS2–MCP-based system used in this study. MS2: bacteriophage MS2 RNA sequence; MCP: MS2 coat protein. (F, G) HEK293T cells transfected with the indicated expression plasmids. Cell lysates were subjected to IP. Immunoprecipitates and TCLs were analyzed by IB. In G, some immunoprecipitates bound to beads were treated with RNase A followed by washing, and the resulting precipitates were analyzed by IB.

To test this, we used three Gag mutants: Gag^14KR^, Gag^4KR,^ and an additional mutant, Gag^2KR^ (two lysines in p6 replaced) (Fig. 3A). Co-immunoprecipitation analysis showed that the Gag-ALIX interaction was reduced in Gag^2KR^ and Gag^4KR^ and nearly abolished in Gag^14KR^ (Fig. 4B). Expression of USP8^C786A^ enhanced the Gag-ALIX interaction compared to wild-type USP8 (Fig. 4C). Similarly, the expression of UbV8.2^ΔGG^ also increased this interaction (Fig. 4D). These results support the notion that Gag ubiquitination promotes ALIX interactions.

Next, we examined whether ALIX bound to vRNA using a bacteriophage MS2-MS2 coat protein (MS2-MCP)-based system (30) (Fig. 4E). ALIX coprecipitated with vRNA-MS2 via MS2-MCP (Fig. 4F, lane 2). The Bro1 domain was responsible for this interaction (lanes 3 and 4), consistent with a previous report suggesting that Bro1 possesses RNA-binding ability (27). RNase A treatment reduced Bro1 co-precipitation, confirming its direct interaction with vRNA (Fig. 4G).

### ALIX enhances Gag-vRNA binding and vRNA packaging

ALIX knockout (KO) cells were used to assess a role of ALIX in Gag-vRNA binding. Gag-vRNA binding decreased to ∼25% of that of the control in ALIX KO cells (Fig. 5A and 5B) without altering intracellular vRNA levels (Fig. S4A). vRNA packaging into virions also decreased (Fig. 5C), whereas the amount of intracellular vRNA unexpectedly increased (Fig. 5D). Gag protein levels in the virions were unaffected (Fig. 5E).

**Figure 5.**
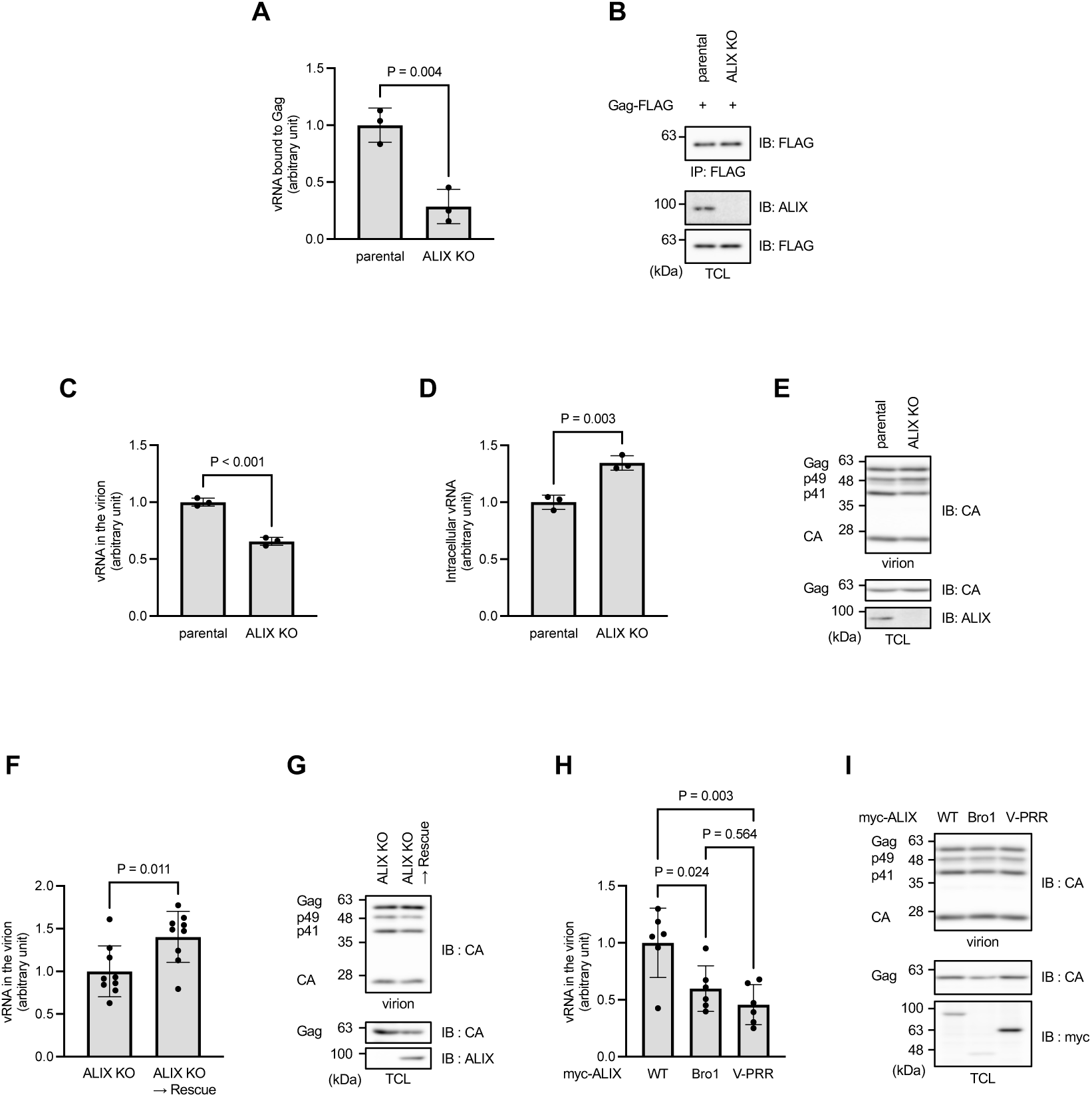
ALIX enhances Gag-vRNA binding and vRNA packaging. (A, B) Parental HEK293T cells and ALIX knockout (KO) cells were transfected with the expression plasmids encoding FLAG-tagged Gag and vRNA. Cell lysates were subjected to IP with anti-FLAG-antibody. vRNA levels in the immunoprecipitates were quantified by RT-qPCR (A). Gag and ALIX levels in immunoprecipitates and TCLs were analyzed by IB (B). (C-E) Lentiviral vectors encoding a luciferase reporter were produced in parental or ALIX KO cells. vRNA levels in the viral fractions (C) and cellular vRNA levels (D) were quantified by RT-qPCR. The amounts of Gag, its processed forms, and ALIX in the viral fractions and TCLs were analyzed by IB (E). (F-I) Lentiviral vectors were produced in ALIX KO cells transfected with the expression plasmids encoding full-length or its truncated forms of ALIX. vRNA levels in the viral fractions were quantified by RT-qPCR (F, H). The amounts of Gag, its processed forms, and ALIX in the viral fractions and TCLs were analyzed by IB (G, I). Graphs show means ± SD from three (A, C, D), nine (F), and six (H) biological replicates.

Re-expression of ALIX restored vRNA packaging (Fig. 5F) without affecting intracellular vRNA or Gag levels (Figs. S4B and 5G). When individual domains (Bro1 or V-PRR) were expressed, the vRNA levels in the virions were significantly lower than those in full-length ALIX (Figs. 5H, 5I, and S4C), suggesting that both domains are necessary for full ALIX activity.

### The USP8-Gag-ALIX axis regulates Gag-vRNA binding

To test whether USP8 and ALIX act via the same axis, we analyzed Gag-vRNA binding under USP8 modulation. In cells expressing Gag and vRNA, co-expression of USP8^C786A^ increased Gag-vRNA binding compared to wild-type USP8 (Fig. 6A and 6B). This enhancement was abolished by the Gag 14KR mutation (Fig. 6C and 6D). Similarly, UbV8.2^ΔGG^ increased Gag-vRNA binding (Fig. 6E and 6F), thereby supporting the role of USP8 as a negative regulator. Importantly, in ALIX KO cells, neither USP8^C786A^ nor UbV8.2^ΔGG^ altered Gag-vRNA binding (Figs. 6G–J, S5A, and S5B), indicating that ALIX mediates the effect of USP8 on Gag-vRNA binding. Collectively, our findings support the hypothesis that USP8 deubiquitinates Gag, thereby reducing its interaction with ALIX and consequently diminishing Gag-vRNA binding (Fig. 6K).

**Figure 6.**
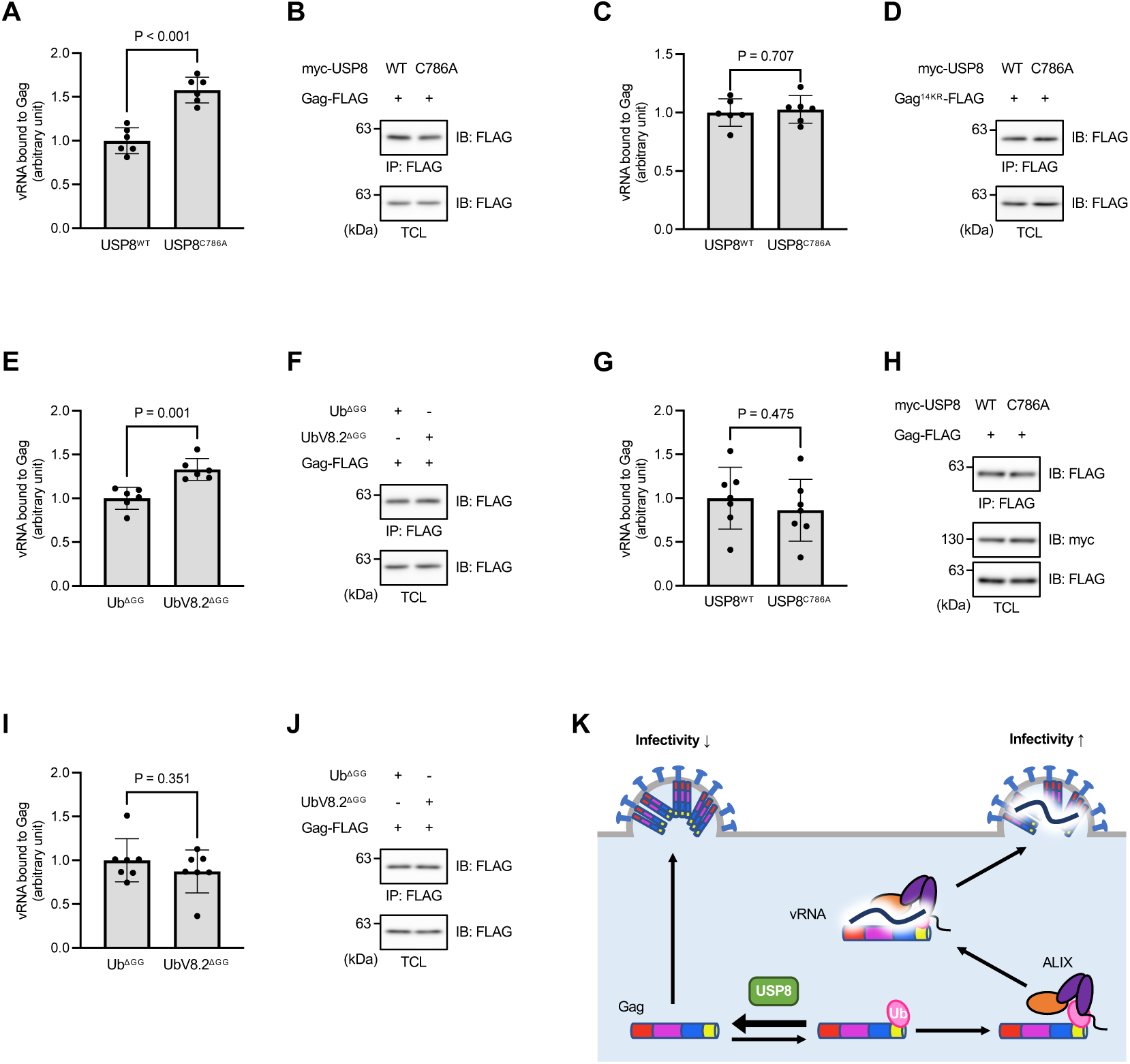
The USP8-Gag-ALIX axis regulates Gag-vRNA binding. (A, B) HEK293T cells were transfected with the expression plasmids encoding FLAG-tagged Gag, vRNA, and either wild-type or C786A USP8. Cell lysates were subjected to IP with anti-FLAG-antibody. vRNA levels in the immunoprecipitates were quantified by RT-qPCR (A). Gag levels in immunoprecipitates and TCLs were analyzed by IB (B). (C, D) Similar experiments were performed as described in A and B, except that a plasmid encoding Gag^14KR^ was used instead of wild-type Gag. (E, F) Similar experiments were performed as described in A and B, except that the indicated ubiquitin-related plasmids were used instead of USP8 expression plasmids. (G, H) Similar experiments were performed as described in A and B, except that ALIX KO cells were used instead of parental HEK293T cells. (I, J) Similar experiments were performed as described in E and F, except that ALIX KO cells were used instead of parental HEK293T cells. (K) Schematic model of the new regulatory mechanism for lentiviral assembly. Graphs show means ± SD from six (A, C, E) and seven (G. I) biological replicates.

## Discussion

USP8 is associated with antiviral response. This effect is mediated by several molecular mechanisms. For example, USP8 deubiquitinates and stabilizes cytidine deaminase APOBEC3G, a key antiviral factor (18). It also stabilizes IFNAR2 by removing ubiquitin chains, thereby enhancing type I interferon (IFN-I)-mediated antiviral signaling (21). Upon stimulation with double-stranded DNA that mimics viral infection, USP8 translocates to stress granules (SGs) and deubiquitinates the SG-associated protein DDX3X, which enhances cGAS-STING signaling and IFN secretion (23). In this study, we propose a new antiviral mechanism in which USP8 deubiquitinates lentiviral Gag, thereby suppressing Gag-vRNA binding and subsequent vRNA packaging into virions. However, it should be noted that USP8 overexpression enhanced viral infectivity independent of its catalytic activity, possibly by acting as a scaffold protein (Fig. 1B). The activity-independent function of this enzyme requires further investigation. Collectively, these findings indicate that USP8 exerts diverse antiviral effects as a host factor. Interestingly, lentiviral HIV-1 infection reduces USP8 expression in human T cells (18, 31, 32), potentially dampening the host antiviral response.

We also identified a new role of Gag ubiquitination in vRNA packaging. Previous studies have reported that Nedd4-1/2 ubiquitinated Gag to promote lentiviral budding (12–16). However, under our experimental conditions, the Gag 14KR mutation did not reduce the amount of Gag in the viral fraction but markedly decreased the vRNA content (Fig. 3), suggesting that Gag ubiquitination is dispensable for budding but essential for vRNA packaging. This discrepancy may stem from the use of Gag constructs lacking binding sites for ALIX or tumor susceptibility gene 101 protein (Tsg101) (12–16), which are key factors involved in budding. The functional consequences of Gag ubiquitination may depend not only on the Gag sequence but also on ubiquitin topology (e.g., monoubiquitin vs. polyubiquitin). In our experiments, Gag was primarily mono-ubiquitinated. In contrast, another group reported that DUB inhibitors induce Gag polyubiquitination, leading to proteasomal degradation and antigen presentation (33). Gag also noncovalently associates with free ubiquitin chains and other ubiquitinated proteins (Fig. 2C). These findings underscore the need for further studies to elucidate the specific roles of different ubiquitin moieties bound to Gag.

ALIX binds to Gag and recruits budding executor ESCRT-III (34). However, as another Gag-binding protein, Tsg101, can also recruit ESCRT-III, and ALIX appears to fulfill this role primarily when the Tsg101-mediated pathway is compromised (35). In this study, we identified a new role for ALIX in lentiviral particle formation. Upon Gag ubiquitination, ALIX binds more strongly to Gag, probably because of the interaction between the ubiquitin moiety on Gag and the ubiquitin-binding site within the ALIX V domain. Moreover, Gag ubiquitination enhances Gag-vRNA binding in an ALIX-dependent manner. A plausible mechanism is that the Gag-ALIX complex contains multiple RNA-binding domains, specifically the Gag NC and ALIX Bro1 domains, which together enable the high-affinity recognition of vRNA. Additionally, ubiquitin-binding may induce a conformational change in ALIX that exposes the Bro1 domain, thereby increasing its accessibility to vRNA. Supporting this speculation, cytoplasmic ALIX proteins adopt a “closed” conformation, in which the PRR interacts with the Bro1 domain (36–38). Consistently, we observed that the isolated Bro1 domain bound to vRNA more strongly than full-length ALIX (Fig. 3G). Further studies are required to elucidate the molecular mechanisms underlying these regulatory effects.

This study highlights their significant potential for both medical and industrial applications. Targeting host factors offers advantages for antiviral therapy development, including a lower risk of drug-resistant viral mutations (39) and broader efficacy across viral subtypes. We showed that lentiviruses hijack the host cell machinery for vRNA packaging, emphasizing the potential of antiviral strategies targeting host proteins, such as USP8 and ALIX. Our findings may enhance the efficiency of industrial-scale lentiviral vector production. A major obstacle in this process is the generation of incomplete vectors lacking transgenes (8). Modulating the USP8-Gag-ALIX axis, for example, by inhibiting USP8, can improve transgene incorporation.

In summary, we propose a new regulatory mechanism for lentiviral assembly, in which Gag ubiquitination facilitates ALIX-mediated vRNA packaging and is counteracted by USP8. These insights will provide a foundation for developing new antiviral strategies and improving lentiviral vector manufacturing.

## Materials and Methods

### Plasmids

The following cDNAs were used for plasmid construction: human *USP8* (40), human *ubiquitin* (40), HIV-1-derived *Gag* (from psPAX2, Addgene #12260), human *ALIX* (cloned from U2OS cell cDNA), twelve tandem copies of the bacteriophage MS2 sequences (from pUB_smHA_KDM5B_MS2; Addgene, #81085), the bacteriophage *MS2 coat protein* (*MCP*) (from pLR5-CAG-MCP-tdiRFP670; kindly provided by Dr. Yuko Sato, Kyushu University), and *NanoLuc* luciferase (Promega) cDNAs. After subcloning, site-directed mutagenesis was performed by PCR using the PrimeSTAR Mutagenesis Basal Kit (Takara) according to the manufacturer’s instructions. Specifically, the following mutants and truncations were generated: a catalytically inactive USP8 mutant (USP8^C786A^), USP8 catalytic domain (amino acids 756-1118), ubiquitin ΔGG mutant (Ub^ΔGG^, in which the C-terminal diglycine motif was deleted), USP8-inhibitory ubiquitin mutant (UbV8.2^ΔGG^, containing 12 point mutations (24) and lacking the C-terminal diglycine motif), Gag ubiquitination site mutants (2KR, 4KR, and 14KR, see Fig. 3), ALIX Bro1 domain (amino acids 1-360), and ALIX V-PRR region (amino acids 361-868).

A second-generation lentiviral vector system was also used (41). For virus production, three lentivirus plasmids were used: psPAX2, which expresses HIV-1-derived Gag and Gag-Pol; pCMV-VSV-G (Addgene #8454), which expresses the vesicular stomatitis virus glycoprotein (VSV-G) as an envelope protein to enable infection of a broad range of mammalian cells; and pCDH-CMV-MCS-EF1-Puro (System Biogenesis), which expresses vRNA encoding the HIV-1 packaging signal (psi) and transgene inserted in the multiple cloning site (MCS). Lentiviral vectors encoding USP8, USP8^C786A^, or NanoLuc were produced using pCDH-CMV-FLAG-USP8-EF1-Puro (40), pCDH-CMV-FLAG-USP8^C786A^-EF1-Puro, or pCDH-CMV-NanoLuc-EF1-Puro plasmids, respectively. Lentiviral vectors with Gag mutations (4KR and 14KR) were generated using psPAX2 harboring 4KR and 14KR mutations.

We constructed the following mammalian expression plasmids: myc-USP8 (wild-type and C786A) (pCMV5-myc vector kindly provided by Dr. Atsushi Nakai, Keio University), HA-ubiquitin (pcDNA3) (40), myc-Ub^ΔGG^ (pCMV5-myc), myc-Ubv8.2^ΔGG^ (pCMV5-myc), FLAG-Ub^ΔGG^ (pFLAG-CMV2, Sigma-Aldrich), FLAG-Ubv8.2^ΔGG^ (pFLAG-CMV2), Gag-FLAG (wild-type, 2KR, 4KR and 14KR) (pCMV-FLAG, kindly provided by Dr. Masanori Iwaki, Osaka University), Gag-HA (wild-type, 2KR, 4KR and 14KR) (pCMV-FLAG with a 2×HA tag and a stop codon inserted between the Gag and FLAG sequences), myc-ALIX (full-length, Bro1 domain, and V-PRR; pCMV-myc), and FLAG-ALIX (pFLAG-CMV2). We also constructed the following bacterial expression plasmids: GST-USP8 catalytic domain (amino acids 756-1118; pGEX, Cytiva), His-Ub^ΔGG^ (pET-15b, Novagen), and His-UbV8.2^ΔGG^ (pET-15b). pCDH-CMV-MCP-FLAG-MS2-EF1-puro was constructed as a dual-expression vector encoding MCP-FLAG and vRNA containing 12 MS2 repeats (vRNA-MS2).

### Cell culture and transfection

HEK293T and HeLa cells were maintained in Dulbecco’s Modified Eagle’s Medium (Nacalai Tesque) supplemented with 10% fetal bovine serum (FBS), 100 U/ml penicillin, and 100 μg/ml streptomycin at 37°C and 5% CO_2_. Plasmid transfection was performed using polyethylenimine (Polysciences) according to the standard protocol. Transfection of ON-TARGETplus Human USP8 siRNA (# J-005203-08-0010; Dharmacon) and a non-targeting siRNA pool (# D-001810-10-20; Dharmacon) was performed using Lipofectamine RNAiMAX (Invitrogen) following the manufacturer’s instructions.

### Generation of ALIX KO cells

ALIX knockout (KO) cells were generated using the CRISPR-Cas9 system. A guide RNA (gRNA) targeting exon 1 of the *ALIX* gene was designed using CRISPRdirect (https://crispr.dbcls.jp) with the sequence, 5’-AGCAGACTTACCCAAGCGGC-3’. gRNA was inserted into the pX459P vector (kindly provided by Dr. Takeshi Shimi, Kanazawa University) and transfected into HEK293T cells using FuGENE 4 K (Promega). Forty-eight hours after transfection, cells were selected with 3 μg/ml puromycin (InvivoGen) for 24 h, followed by an additional 48 h with 5 μg/ml puromycin. Single-cell clones were isolated using limiting dilution. Genomic DNA of each clone was extracted by lysing the cells with lysis buffer (0.1 mM Tris-HCl, pH 8.0, 200 mM NaCl, 5 mM EDTA, 0.4% SDS, 0.2 mg/ml Proteinase K) at 55°C for 40 min. After isopropanol precipitation, DNA was resuspended in TE buffer (10 mM Tris-HCl, pH 7.4, 1 mM EDTA) and subjected to PCR to amplify the region flanking the gRNA target site, using PrimeSTAR Max DNA Polymerase (Takara) with the following primers: 5’-TCTCGGTGCAGCTGAAAAAG-3’ and 5’-GAAAAATGGGGCGAAAGGGG-3’. PCR products were purified by agarose gel electrophoresis, followed by gel extraction of the target bands using a Gel/PCR Extraction Kit (Nippon Genetics). Sanger sequencing was performed using the primer 5’-ATCAGTACAGGGCAGACTGC-3’. Clones with frameshift mutations in both ALIX alleles were used in subsequent experiments.

### Lentiviral production, purification, and infection

For lentiviral production, HEK293T and ALIX KO cells were transfected with pCDH-CMV-MCS-EF1-Puro, which encodes the transgene, along with psPAX2 and pCMV-VSV-G. Twenty-four hours after transfection, the medium was replaced, and after an additional 24 h, the medium was collected and filtered. For purification, the filtered medium was layered onto PBS containing 20% glucose in a 13PA tube (Himac) and centrifuged at 25,000 rpm at 4°C for 90 min using a CP80β ultracentrifuge (Eppendorf) equipped with a swinging bucket rotor (P40ST, Eppendorf). The pellet was washed with PBS and centrifuged again at 25,000 rpm at 4°C for 15 min. The resulting pellet was used as the viral fraction for subsequent analysis. For infection, the filtered viral medium was diluted (1:100) and added to HeLa cells in the presence of 10 μg/mL polybrene (Nacalai Tesque).

### Measurement of viral infectivity

Infectivity was quantified using the crystal violet assay. Specifically, 48 h after infection, HeLa cells were cultured in the presence of 1.6 μg/mL puromycin for an additional 48 h. The surviving cells were washed twice with PBS and incubated with crystal violet solution (0.5% crystal violet, 20% methanol) at 20 °C for 20 min. After three washes with distilled water and air drying at room temperature for 24 h, methanol was added, and the dye was gently extracted by shaking for 20 min. The extracts were diluted three-fold, and the absorbance was measured at 570 nm using a Varioskan LUX (Thermo Scientific).

The infectivity of lentiviruses carrying the NanoLuc sequence was determined using a luciferase assay. Briefly, 48 h post-infection, the Nano-Glo Luciferase Assay System (Promega) and Reporter Lysis 5×Buffer (Promega) were used according to the manufacturer’s protocols.

### Cell lysis and immunoprecipitation

Forty-eight hours after transfection, cells were lysed with ice-cold lysis buffer (50 mM Tris-HCl, pH 7.4, 150 mM NaCl, 50 mM NaF, 1 mM EDTA, 1 mM EGTA, 1% Triton X-100, 4.0 μg/mL leupeptin, 3.3 μg/mL pepstatin, 2.5 μg/mL aprotinin). To prevent the deubiquitination of Gag, 2 mM N-ethylmaleimide was added to the lysis buffer. The lysates were then centrifuged at 13,500 rpm at 4°C for 10 min, and the supernatants were incubated with either anti-FLAG M2 affinity gel (Sigma-Aldrich) or anti-DYKDDDDK tag antibody-conjugated beads (Wako) with rotation at 4°C for 2 h. The beads were washed five times with lysis buffer, and FLAG-tagged proteins were eluted by incubating the beads with lysis buffer containing 200 μg/mL FLAG peptide (Sigma-Aldrich) at 4°C for 30 min.

In the crosslink IP experiments shown in Fig. 2A, before cell lysis, 0.1% formaldehyde was added to the culture medium and incubated at room temperature for 10 min. Subsequently, 133 mM glycine was added, and the mixture was incubated for 5 min. The cells were lysed as described above, followed by sonication (BRANSON, output 1, constant for 15 s). The lysates were centrifuged at 13,500 rpm at 4°C for 10 min, and the supernatants were used for IP.

For the denaturing procedure to remove non-covalently bound ubiquitin from Gag in Fig. 2C, the lysates were mixed with 2% SDS and then incubated at 99°C for 5 min. The samples were then diluted 20-fold with lysis buffer and incubated at 4°C for 30 min before IP.

For RNase A treatment (Fig. 4G), the precipitated beads were washed once with TE buffer and then incubated with TE buffer containing 50 μg/ml RNase A (QIAGEN) at 37°C for 1 h. Subsequently, the beads were washed, and the proteins were eluted as described above.

### Immunoblotting

Cell lysates, immunoprecipitates, and viral fractions were incubated in SDS-PAGE sample buffer (100 mM Tris-HCl, pH 6.8, 2% SDS, 2.5% 2-mercaptoethanol, 5% glycerol, and 0.05% bromophenol blue) at 98°C for 5 min. Immunoblotting (IB) was performed according to standard protocols. The following primary antibodies were used: anti-FLAG antibody (clone 1E6, Wako, 0.1– 0.5 μg/mL), anti-USP8 rabbit antiserum (1:200 dilution) (42), anti-HA antibody (clone 3F10, Roche, 0.1 μg/mL), anti-myc antibodies (clone 9E10, Wako, 1.0 μg/mL; and clone 1A5A2, Proteintech, 0.5 μg/mL), anti-CA antibodies (clone 39/5.4A, Abcam, 1:1,000; and clone 24-4, Santa Cruz Biotechnology, 0.4 μg/mL), and anti-ALIX antibody (clone 3A9, Santa Cruz Biotechnology, 1.0 μg/mL). The following horseradish peroxidase-conjugated secondary antibodies were used: anti-mouse IgG antibody (GE Healthcare, 1:40,000), anti-rabbit IgG antibody (GE Healthcare, 1:40,000), and anti-rat IgG antibody (GE Healthcare, 1:20,000). Chemiluminescent signals were detected using ECL Prime Western Blotting Detection Reagents (GE Healthcare) and SuperSignal Femto (Thermo Scientific) and visualized using an ImageQuant LAS 4000 mini system (GE Healthcare).

### Immunofluorescence staining

Forty-eight hours after transfection, HeLa cells cultured on coverslips were fixed with 4% paraformaldehyde at room temperature for 10 min. The cells were permeabilized with PBS containing 0.2% Triton X-100 for 10 min, followed by blocking with PBS supplemented with 5% FBS for 45 min. The following primary antibodies were applied overnight at 4°C: rabbit anti-FLAG antibody (Sigma, 1.0 μg/mL), and mouse anti-myc antibody (clone 9E10, Wako, 1.0 μg/mL). After washing, cells were incubated with the following secondary antibodies at room temperature for 30 min: Alexa Fluor 488-conjugated anti-rabbit IgG (Life Technologies, 1:500) and Alexa Fluor 594-conjugated anti-mouse IgG (Life Technologies, 1:500). The coverslips were mounted on glass slides using Fluoroshield mounting medium (ImmunoBioScience). Images were acquired using a laser scanning confocal microscope (LSM 780; Carl Zeiss).

### Protein purification

GST-tagged USP8 (amino acids 756–1118) was purified, as previously described (40). His-tagged Ub^ΔGG^ and UbV8.2^ΔGG^ were expressed in *E. coli* Rosetta cells. Cells were lysed in ice-cold PBS containing 1% Triton X-100 and protease inhibitors (4.0 μg/mL leupeptin, 3.3 μg/mL pepstatin, and 2.5 μg/mL aprotinin), followed by sonication (BRANSON, output 3, 30% duty cycle, for 5 min). The lysates were centrifuged at 13,500 rpm at 4°C for 10 min. TALON Metal Affinity Resin (Clontech Laboratories) was added to the supernatant and incubated with rotation at 4°C for 1 h. The beads were washed with PBS containing 10 mM imidazole and 1% Triton X-100, and His-tagged proteins were eluted with PBS containing 150 mM imidazole and 1% Triton X-100 at 4°C for 10 min. Imidazole was removed by ultrafiltration, and the protein concentration was determined using the Bradford protein assay.

### In vitro deubiquitination assay

The in vitro deubiquitination assay was performed as previously described (40). Briefly, GST-USP8 (amino acids 756–1118) was mixed with 1 μM ubiquitin-MCA (Peptide Institute), 10 mM DTT, and either 5 μM His-Ub^ΔGG^ or His-UbV8.2^ΔGG^ in Tris-buffered saline (TBS; 20 mM Tris-HCl pH 7.4, 150 mM NaCl). The mixture was incubated at 37°C in a Varioskan LUX plate reader (Thermo Scientific), and the fluorescence intensity of MCA released from ubiquitin was measured at an excitation wavelength of 345 nm and an emission wavelength of 445 nm at 10-second intervals.

### RT-qPCR for quantifying vRNA levels

RNA was isolated from cell lysates, immunoprecipitates, and viral fractions using either the FastGene RNA Basic Kit (Japan Genetics) or Sepasol-RNA I Super G reagent (Nacalai Tesque). Reverse transcription was performed using ReverTra Ace qPCR RT Master Mix with gDNA Remover (TOYOBO) according to the manufacturer’s protocol. Quantitative PCR was conducted on a StepOnePlus Real-Time PCR System (Applied Biosystems) using the THUNDERBIRD Next qPCR Mix (TOYOBO) in accordance with the manufacturer’s instructions. The NanoLuc sequence inserted into the vRNA was specifically amplified using the following primers: Forward, 5’-TTACGCCGAACATGATCGAC-3’; Reverse, 5’-TTTTGTTGCCGTTCCACAGG-3 ’.

### Statistical analysis

Statistical analyses were performed using GraphPad Prism version 10 (DotMatics). Data are presented as mean ± standard deviation (SD). An unpaired two-tailed Student’s t-test was used for comparisons between two groups, whereas one-way analysis of variance (ANOVA) followed by Tukey’s post hoc test was used for comparisons among multiple groups. Statistical significance was set at p < 0.05.

## Acknowledgments

We thank Drs. Akira Kato and Naonobu Fujita (Institute of Science Tokyo) for helpful discussions and Drs. Yuko Sato (Kyushu University) and Takeshi Shimi (Kanazawa University) for the generous donation of plasmids. We also acknowledge the Biomaterials Analysis Division at the Institute of Science Tokyo for assistance with the DNA sequencing analysis. We thank Editage and ChatGPT for their assistance in the English language editing process. This work was supported by JSPS KAKENHI Grant Numbers JP19H05289 and JP21H00276, AMED Grant Numbers JP22ym0126806j0001 and JP23ym0126806j0002 (to T.F.), and JSPS KAKENHI Grant Number JP23K21308 (to M.K.).

## References

1. A. Rein, RNA Packaging in HIV. Trends Microbiol. 27, 715–723 (2019).

2. E. Mailler, et al., The Life-Cycle of the HIV-1 Gag–RNA Complex. Viruses 8, 248 (2016).

3. M. Comas-Garcia, S. Davis, A. Rein, On the Selective Packaging of Genomic RNA by HIV-1. Viruses 8, 246 (2016).

4. G. Lerner, N. Weaver, B. Anokhin, P. Spearman, Advances in HIV-1 Assembly. Viruses 14, 478 (2022).

5. E. O. Freed, HIV-1 assembly, release and maturation. Nat. Rev. Microbiol. 13, 484–496 (2015).

6. J. R. Lingappa, J. C. Reed, M. Tanaka, K. Chutiraka, B. A. Robinson, How HIV-1 Gag assembles in cells: Putting together pieces of the puzzle. Virus Res. 193, 89–107 (2014).

7. S. J. Rulli, et al., Selective and nonselective packaging of cellular RNAs in retrovirus particles. J. Virol. 81, 6623–6631 (2007).

8. C. Perry, A. C. M. E. Rayat, Lentiviral Vector Bioprocessing. Viruses 13, 268 (2021).

9. D. E. Ott, et al., Ubiquitin is covalently attached to the p6Gag proteins of human immunodeficiency virus type 1 and simian immunodeficiency virus and to the p12Gag protein of Moloney murine leukemia virus. J. Virol. 72, 2962–2968 (1998).

10. D. E. Ott, L. V. Coren, E. N. Chertova, T. D. Gagliardi, U. Schubert, Ubiquitination of HIV-1 and MuLV Gag. Virology 278, 111–121 (2000).

11. E. Gottwein, S. Jäger, A. Habermann, H.-G. Kräusslich, Cumulative Mutations of Ubiquitin Acceptor Sites in Human Immunodeficiency Virus Type 1 Gag Cause a Late Budding Defect. J. Virol. 80, 6267–6275 (2006).

12. H.-Y. Chung, et al., NEDD4L Overexpression Rescues the Release and Infectivity of Human Immunodeficiency Virus Type 1 Constructs Lacking PTAP and YPXL Late Domains. J. Virol. 82, 4884–4897 (2008).

13. P. Sette, J. A. Jadwin, V. Dussupt, N. F. Bello, F. Bouamr, The ESCRT-associated protein Alix recruits the ubiquitin ligase Nedd4-1 to facilitate HIV-1 release through the LYPXnL L domain motif. J. Virol. 84, 8181–8192 (2010).

14. A. Joshi, U. Munshi, S. D. Ablan, K. Nagashima, E. O. Freed, Functional replacement of a retroviral late domain by ubiquitin fusion. Traffic Cph. Den. 9, 1972–1983 (2008).

15. Y. Usami, S. Popov, E. Popova, H. G. Göttlinger, Efficient and Specific Rescue of Human Immunodeficiency Virus Type 1 Budding Defects by a Nedd4-Like Ubiquitin Ligase. J. Virol. 82, 4898–4907 (2008).

16. E. R. Weiss, et al., Rescue of HIV-1 Release by Targeting Widely Divergent NEDD4-Type Ubiquitin Ligases and Isolated Catalytic HECT Domains to Gag. PLoS Pathog. 6, e1001107 (2010).

17. A. Dufner, K.-P. Knobeloch, Ubiquitin-specific protease 8 (USP8/UBPy): a prototypic multidomain deubiquitinating enzyme with pleiotropic functions. Biochem. Soc. Trans. 47, 1867–1879 (2019).

18. W. Gao, et al., Specific Deubiquitinating Enzymes Promote Host Restriction Factors Against HIV/SIV Viruses. Front. Immunol. 12, 740713 (2021).

19. W. Xiong, et al., USP8 inhibition reshapes an inflamed tumor microenvironment that potentiates the immunotherapy. Nat. Commun. 13, 1700 (2022).

20. Y. Miao, et al., Spatiotemporal recruitment of the ubiquitin-specific protease USP8 directs endosome maturation. eLife 13, RP96353 (2024).

21. J. Guo, H. Zheng, S. Xiong, SENP6 restricts the IFN-I-induced signaling pathway and antiviral activity by deSUMOylating USP8. Cell. Mol. Immunol. 21, 892–904 (2024).

22. A. Endo, et al., USP8 prevents aberrant NF-κB and Nrf2 activation by counteracting ubiquitin signals from endosomes. J. Cell Biol. 223, e202306013 (2024).

23. X. Zhang, et al., Stress granule-localized USP8 potentiates cGAS-mediated type I interferonopathies through deubiquitination of DDX3X. Cell Rep. 43, 114248 (2024).

24. A. Ernst, et al., A strategy for modulation of enzymes in the ubiquitin system. Science 339, 590–595 (2013).

25. S. Lee, A. Joshi, K. Nagashima, E. O. Freed, J. H. Hurley, Structural basis for viral late-domain binding to Alix. Nat. Struct. Mol. Biol. 14, 194–199 (2007).

26. S. Popov, E. Popova, M. Inoue, H. G. Göttlinger, Human immunodeficiency virus type 1 Gag engages the Bro1 domain of ALIX/AIP1 through the nucleocapsid. J. Virol. 82, 1389– 1398 (2008).

27. P. Sette, V. Dussupt, F. Bouamr, Identification of the HIV-1 NC Binding Interface in Alix Bro1 Reveals a Role for RNA. J. Virol. 86, 11608–11615 (2012).

28. D. P. Dowlatshahi, et al., ALIX is a Lys63-specific polyubiquitin binding protein that functions in retrovirus budding. Dev. Cell 23, 1247–1254 (2012).

29. T. Keren-Kaplan, et al., Structure-based in silico identification of ubiquitin-binding domains provides insights into the ALIX-V:ubiquitin complex and retrovirus budding. EMBO J. 32, 538–551 (2013).

30. E. Bertrand, et al., Localization of ASH1 mRNA particles in living yeast. Mol. Cell 2, 437– 445 (1998).

31. N. J. Matheson, et al., Cell Surface Proteomic Map of HIV Infection Reveals Antagonism of Amino Acid Metabolism by Vpu and Nef. Cell Host Microbe 18, 409–423 (2015).

32. E. J. Greenwood, et al., Temporal proteomic analysis of HIV infection reveals remodelling of the host phosphoproteome by lentiviral Vif variants. eLife 5, e18296 (2016).

33. C. Setz, et al., Inhibitors of Deubiquitinating Enzymes Block HIV-1 Replication and Augment the Presentation of Gag-Derived MHC-I Epitopes. Viruses 9, 222 (2017).

34. K. M. Rose, V. M. Hirsch, F. Bouamr, Budding of a Retrovirus: Some Assemblies Required. Viruses 12, 1188 (2020).

35. Y. Usami, S. Popov, H. G. Göttlinger, Potent rescue of human immunodeficiency virus type 1 late domain mutants by ALIX/AIP1 depends on its CHMP4 binding site. J. Virol. 81, 6614– 6622 (2007).

36. X. Zhou, J. Si, J. Corvera, G. E. Gallick, J. Kuang, Decoding the intrinsic mechanism that prohibits ALIX interaction with ESCRT and viral proteins. Biochem. J. 432, 525–534 (2010).

37. Q. Zhai, et al., Activation of the retroviral budding factor ALIX. J. Virol. 85, 9222–9226 (2011).

38. R. D. Elias, B. Ramaraju, L. Deshmukh, Mechanistic roles of tyrosine phosphorylation in reversible amyloids, autoinhibition, and endosomal membrane association of ALIX. J. Biol. Chem. 297, 101328 (2021).

39. A. Zumla, et al., Host-directed therapies for infectious diseases: current status, recent progress, and future prospects. Lancet Infect. Dis. 16, e47–63 (2016).

40. K. Kakihara, et al., Molecular basis of ubiquitin-specific protease 8 autoinhibition by the WW-like domain. Commun. Biol. 4, 1272 (2021).

41. T. Sakuma, M. A. Barry, Y. Ikeda, Lentiviral vectors: basic to translational. Biochem. J. 443, 603–618 (2012).

42. M. Kato, K. Miyazawa, N. Kitamura, A deubiquitinating enzyme UBPY interacts with the Src homology 3 domain of Hrs-binding protein via a novel binding motif PX(V/I)(D/N)RXXKP. J. Biol. Chem. 275, 37481–37487 (2000).

