## Supplementary figures for "Gag ubiquitination facilitates ALIX-mediated lentiviral RNA packaging, counteracted by USP8"

b. Cell biology center, Institute of integrated research, Institute of Science Tokyo, 4259 Nagatsuta, Midori-ku, Yokohama, 226-8501, Japan.

1. equal contribution[1]

*Masayuki Komada, Toshiaki Fukushima

**This file includes:**

Figures S1 to S5

Legends for figure S1 to S5

1

**Supplementary Figure S1**

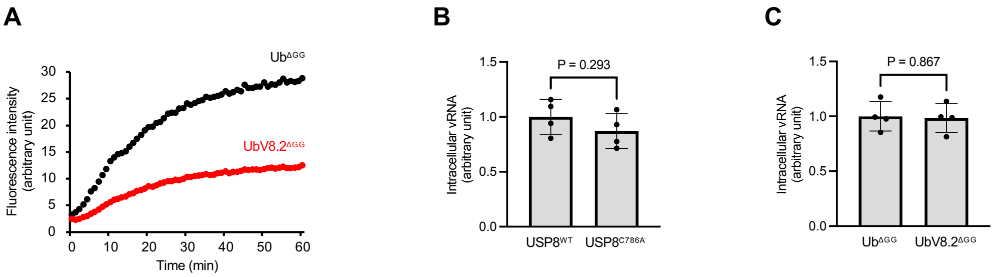


**Figure S1. Data related to Figure 1.**

(A) The deubiquitinating activity of GST-tagged USP8 catalytic domain (amino acids 756-1118) was assessed in the presence of His-tagged ubiquitin mutants (Ub$^{ΔGG}$ or UbV8.2$^{ΔGG}$). Reactions were performed using ubiquitin-MCA as a substrate, and the fluorescence intensity of released MCA was measured. The result confirmed that UbV8.2$^{ΔGG}$ inhibits USP8 activity.

(B, C) Cellular vRNA levels in HEK293T cells used in Figures 1B and 1C were quantified by RT-qPCR.

**Supplementary Figure S2**

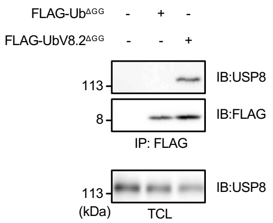


**Figure S2. Data related to Figure 2.**

HEK293T cells were transfected with the indicated plasmids. Cell lysates were subjected to immunoprecipitation. Immunoprecipitates and total cell lysates were analyzed using immunoblotting. The result confirmed that UbV8.2$^{\Delta GG}$ binds to USP8 more strongly than Ub$^{\Delta GG}$.

**Supplementary Figure S3**

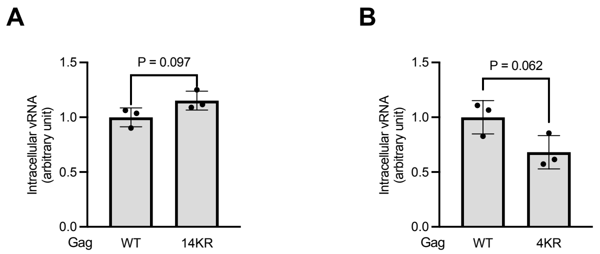


**Figure S3. Data related to Figure 3.**

The cellular vRNA levels in HEK293T cells used in Figures 3B and 3C were quantified by RT-qPCR.

**Supplementary Figure S4**

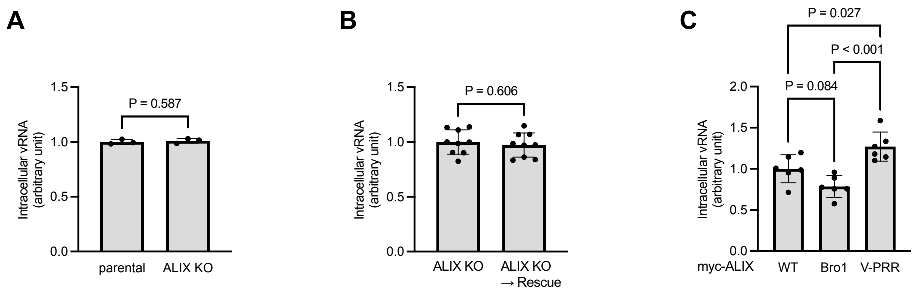


**Figure S4. Data related to Figure 5.**

The cellular vRNA levels in HEK293T cells used in Figures 5A, 5F, and 5H were quantified by RT-qPCR.

**Supplementary Figure S5**

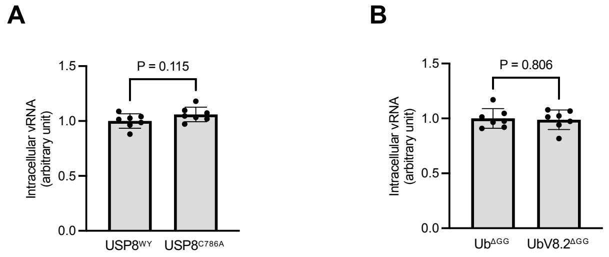


**Figure S5. Data related to Figure 6.**

The cellular vRNA levels in HEK293T cells used in Figures 6G and 6I were quantified by RT-qPCR.
